# PROTFIG: one-command, publication-quality biomolecular figures

**DOI:** 10.64898/2026.10.20.760606

**Authors:** L. América Chi

**Affiliations:** Department of Physics, University of Idaho, Moscow, ID, 83844, USA

**Keywords:** molecular visualization, scientific figures, PyMOL, reproducibility, structural biology, protein–ligand interactions

## Abstract

Molecular structures contain rich information about biomolecular interactions, but translating this information into clear, scientifically informative figures remains a largely manual process. Producing publication-quality illustrations requires decisions about molecular representations, colors, structural features, camera orientation, and annotations, often involving extensive user interaction, expertise and limiting reproducibility. Here, we introduce PROTFIG, an open-source, PyMOL-based command-line tool that transforms biomolecular structures into scientifically informative, reproducible, and editable figures with minimal manual intervention. Given a structure file, PDB identifier or PyMOL scene, PROTFIG automatically identifies relevant molecular components and structural features, selects appropriate visual representations, and generates publication-quality figures using sixteen customizable presets. These cover a broad range of structural contexts, including protein–ligand and protein–peptide interactions, binding pockets, electrostatic surfaces, protein–protein interfaces, oligomeric assemblies, and membrane proteins. Figures can be customized through command-line options or TOML configuration files, while saved PyMOL sessions preserve full editability. We demonstrate the versatility of PROTFIG using representative structures relevant to drug discovery and public health, including GLP-1 receptor agonists, targeted protein degraders, and H5N1 influenza hemagglutinin. By combining automated structural interpretation with reproducible visualization workflows, PROTFIG makes the generation of scientifically informative molecular figures more accessible, consistent, and efficient.

## 1 Introduction

Molecular figures are central to communicating structural findings, translating three-dimensional coordinates into visual representations of biomolecular architecture and interactions. Powerful visualization tools such as PyMOL [1], UCSF ChimeraX [2, 3], and Mol* [4], alongside illustration tools such as Illustrate [5] and Molecular Nodes [6], offer extensive flexibility in how molecular structures are represented. In practice, however, producing a publication-quality molecular figure requires numerous decisions: which chains and residues to display, which representations and colors to use, how to define binding sites and highlight molecular interactions, and how to position the camera, annotations, and lighting. These decisions are rarely documented, making figures difficult to reproduce, maintain consistently across a study, or update when structures are refined or revisions are requested. Scripting can improve reproducibility and consistency, but requires familiarity with the visualization software’s command language.

PROTFIG addresses this gap by making scientifically informative molecular figures easier to generate, reproduce, and modify. Its design follows four principles: (i) *minimal intervention*: a single command transforms a molecular structure into a publication-quality figure; (ii) *scientifically informative presets*: predefined figure designs automatically identify and highlight relevant structural features; (iii) *full editability*: automated decisions can be overridden, and each figure includes an editable PyMOL session; and (iv) *reproducibility*: each figure records the command used to generate it, while complete figure sets can be regenerated from a single configuration file.

## 2 Design and implementation

PROTFIG is written in Python and runs PyMOL headless, following the workflow illustrated in Figure 1a. A typical call is

**Figure 1:**
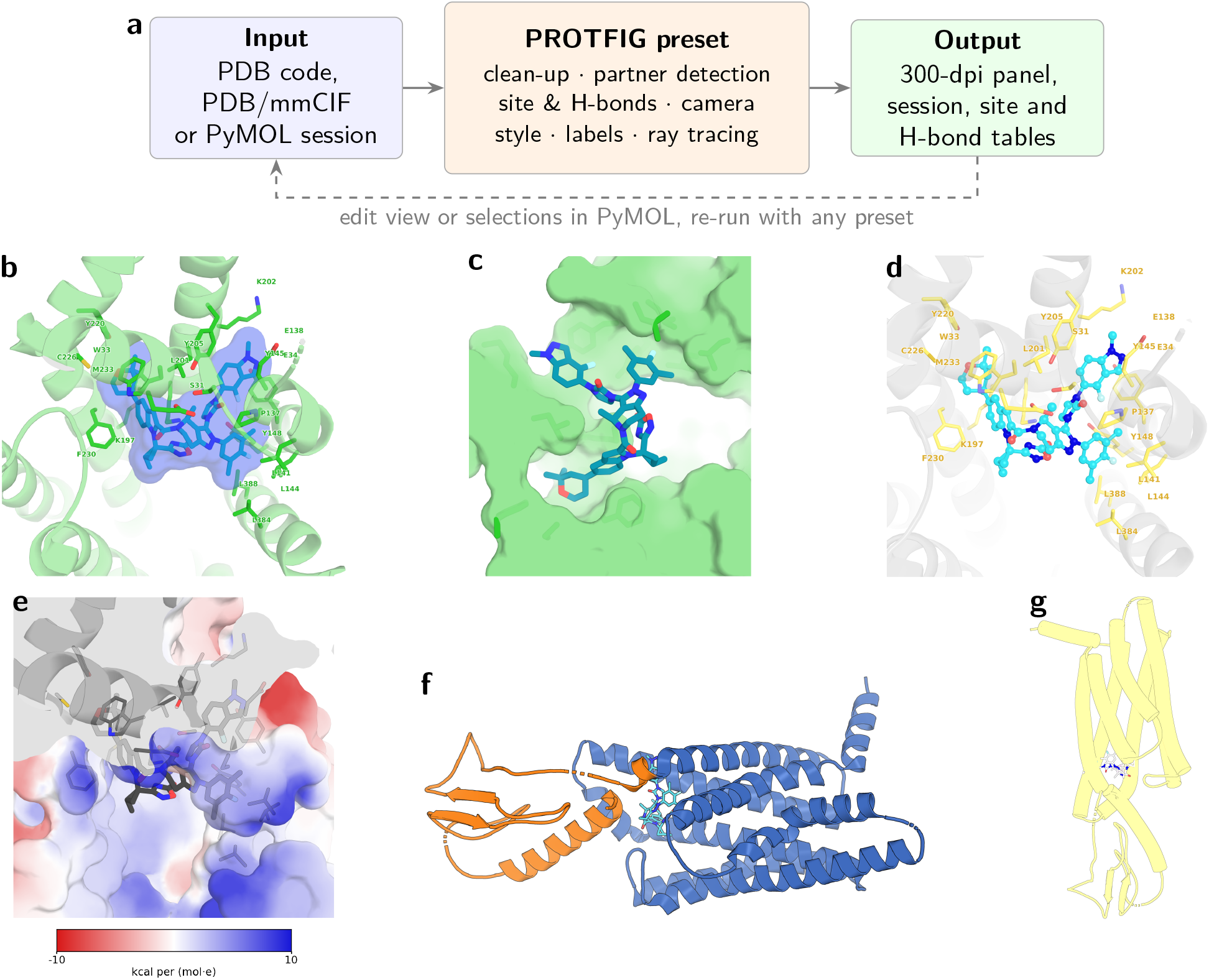
One input, many styles. (**a**) PROTFIG workflow. A single command turns a structure into a finished panel; the saved PyMOL session can be edited and fed back. (**b–g**) The GLP-1 receptor bound to the oral small-molecule agonist orforglipron (LY3502970; pdb `6XOX` [11]) rendered with six presets from the same input, each with one command (`–protfig 6XOX.cif –chains R –preset <name>`): `–site` (b), `–pocket` (c), `–site_faded` (d), `–electrostatic` (e), `–outline` with the extracellular domain highlighted (f, `––highlight “resi 1-140”`) and `–illustrative` (g). The ligand, binding site, hydrogen bonds, camera and labels were determined automatically.

~~~
protfig 6XOX --preset site --chains R
~~~

which retrieves the structure and generates a binding-site figure, automatically highlighting the ligand and surrounding residues (Figure 1b). The resulting panel is saved alongside an editable PyMOL session and annotation files.

### Input and clean-up

Inputs are PDB/mmCIF files, PDB codes or PyMOL sessions. Hydrogens, alternate conformations, crystallographic waters, common crystallization additives (e.g. DMSO, glycerol, sulfate) and the solvent and ions of molecular dynamics (MD) systems are removed; lipids are kept only for the membrane preset.

### Binding partner

Site- and interface-centred presets revolve around a binding partner. Unless specified with `––partner, PROTFIG` automatically selects the largest eligible non-polymeric organic ligand, excluding common additives, lipids, and glycans. If no eligible ligand is found, it selects the smallest polymer chain, classifying it as a peptide (≤50 residues) or protein (*>*50 residues). Presets adapt to the partner type: small molecules are rendered as sticks with an optional translucent molecular surface, whereas peptides and proteins are represented as cartoons with interface side chains and labels on both partners. For large interfaces, labels are restricted to residues involved in hydrogen bonds.

### Binding site and contacts

The binding site comprises receptor residues with at least one heavy atom within a user-defined distance of the partner (`––site_cutoff`, default 4 Å), with hydrogen-bonded residues always included. For protein partners, the same cutoff identifies interacting residues on the partner side of the interface, whereas peptide partners are displayed in full. The `–binding_surface` preset uses a larger cutoff (`––pocket_cutoff`, default 8 Å) to highlight a continuous binding region on the receptor surface rather than only direct-contact residues. Hydrogen bonds are identified as N/O donor–acceptor pairs satisfying PyMOL’s geometric criteria; The same set of hydrogen bonds is used for both the figure and the accompanying text file, ensuring consistency.

### Automatic camera

Each preset automatically orients the camera to emphasize the features of interest. Site views (`–site, –pocket, –site_faded`) face the binding pocket while minimizing ligand foreshortening, making interactions and surrounding residues clearly visible (Figs. 1b–e, 3b–d, and 4c,d). Overview views (`–overview, –outline, –illustrative`) highlight the overall molecular architecture by aligning the shortest principal axis with the line of sight (Figs. 1f,g, 2b, 3a, and 4a). Protein–protein interfaces (`–interface_closeup`) are viewed edge-on, with the partner above the receptor and the interface extending horizontally (Fig. 4b). Binding-footprint views (`–binding_surface`) face the binding region on the receptor surface, switching to a side view when the pocket lies near one end of the receptor’s long axis, as in class B GPCR–peptide complexes (Fig. 2c). Membrane views (`–membrane`) orient the bilayer normal vertically (Fig. 2a). Users can adjust the camera orientation (`––roll, ––turn`), control visibility through clipping (`––clip`), or reuse a saved view or a manually edited PyMOL session.

**Figure 2:**
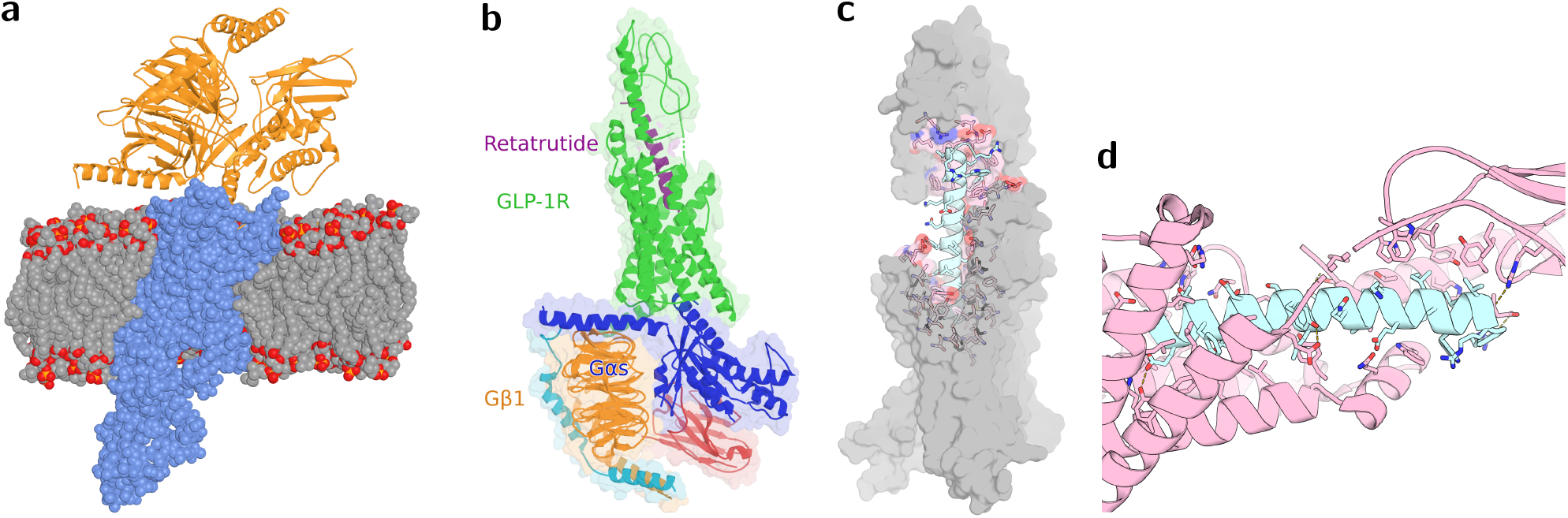
Incretin receptor complexes. (**a**) GLP-1R (blue spheres) in complex with G_s_ and Nb35 (orange cartoon) embedded in a DPPC bilayer (`–membrane`); the cryo-EM complex pdb `6XOX` [11] was superposed onto its MemProtMD simulation [9] (C_*α*_ RMSD 0.20 Å). (**b**) Retatrutide-bound GLP-1R–G_s_ complex (pdb `8YW3` [13]; `–overview` with chain names). (**c, d**) Semaglutide bound to GLP-1R (pdb 7KI0 [14]), detected as a peptide partner: binding footprint on the receptor surface (c, `–binding_surface`) and outlined interface with hydrogen bonds in yellow (d, `–interface_closeup`).

### Labels

Residue labels are anchored at side-chain tips, displaced away from the interaction partner, adjusted to avoid overlap in screen space, and kept within the image boundaries. Labels are rendered as a separate layer and composited over the molecular scene with a thin halo, preventing ray-tracing artifacts in which labels are obscured by translucent cartoons or surfaces. The labels are also exported as an editable SVG layer, alongside the unlabeled molecular image, allowing independent post-processing in vector or raster graphics software.

### Surface properties

Electrostatic surfaces use a Coulomb potential with a distance-dependent dielectric (*ε* = 4*r*), evaluated on a grid from formal charges and mapped onto the surface (±10 kcal mol^−1^ e^−1^ by default). Lipophilicity surfaces use a molecular lipophilicity potential with atom-type contributions and exponential decay [7], with the color range set from the values actually displayed on the surface. Both methods provide fast approximations without requiring external software. More rigorous Poisson– Boltzmann calculations (e.g., APBS [8]) are preferable for quantitative electrostatic analysis. Color bars are exported as editable SVG files.

### Customization and the PyMOL round trip

All visual choices are exposed as options: chain palettes and per-chain colors, partner/receptor/site/hydrogen-bond colors, background, outlines, transparencies, label size, font, color and format, and image size and resolution. Every panel is accompanied by its PyMOL session (`–.pse`), which contains the cleaned structure and editable selections named `–partner` and `–site`. Users may reorient the scene or redefine any selection in PyMOL, save the session and pass it back to PROTFIG, with the same or another preset; the saved view, selections and chain colors are honored.

### Reproducible figure sets

A TOML configuration file defines multiple panels using the same options as the command-line interface. The command `–PROTFIG –config figures.toml –jobs 8` renders them in parallel, saving each panel in a separate directory (`–figures/<preset>/<name>/`) along with metadata recording the command, input structure, selected partner, chains, and generated files. The `––draft` option produces low-resolution previews for rapid inspection. An auxiliary tool allows experimental complexes to be placed within simulated lipid bilayers (e.g., from MemProtMD [9] or CHARMM-GUI [10]) through structural superposition, enabling membrane figures to display complete assemblies. Simulation frames already containing both the protein and bilayer can also be used directly.

## 3 Applications

We applied PROTFIG to structures at the current frontier of drug discovery and public health. All panels in Figures 1–4 were generated with default settings except for the options stated in the legends, and assembled without manual editing.

### Incretin therapeutics

GLP-1 receptor (GLP-1R) agonists have transformed the treatment of obesity and type 2 diabetes. In April 2026, the US FDA approved orforglipron, the first oral small-molecule GLP-1R agonist [12]. Using the orforglipron-bound GLP-1R structure [11], PROTFIG generates six distinct representations from the same input, highlighting the binding site, molecular interactions, electrostatic properties, and overall architecture (Fig. 1b–g). The versatility of PROTFIG is further illustrated with incretin receptor complexes (Fig. 2). These include the GLP-1R–G_s_ signaling complex embedded in a lipid bilayer by structural superposition onto a MemProtMD simulation [9] (Fig. 2a); the triple agonist retatrutide bound to GLP-1R–G_s_ [13] (Fig. 2b); and the peptide drug semaglutide bound to GLP-1R [14] (Fig. 2c,d). For the latter, PROTFIG automatically identifies the peptide partner and generates complementary representations of the binding footprint and molecular interface.

**Table 1.** PROTFIG presets. “Partner” indicates the binding-partner types each preset is designed for; presets without a required partner also work on isolated proteins.

| Preset | Partner | Figure produced |
| --- | --- | --- |
| -overview | optional | Assembly as chain-colored cartoon within a translucent surface; chain names |
| -site | ligand, peptide, protein | Binding-site close-up: site residues, partner, hydrogen bonds, labels |
| -pocket | ligand, peptide | Partner lying in its groove: surface cut behind the partner |
| -site_faded | ligand, peptide, protein | Site over a ghosted cartoon, ball-and-stick ligand, distances; mutations, reference ligand |
| -interface_closeup | ligand, peptide, protein | Outlined close-up of an interface with hydrogen bonds |
| -binding_surface | ligand, peptide, protein | Binding footprint on the receptor surface |
| -outline | optional | Outlined cartoon with highlighted region and key residues |
| -illustrative | optional | Flat-shaded, outlined cylinders or surfaces |
| -pastel | optional | Soft-lit pastel cartoon with N→C gradient |
| -oligomer | none | Outlined surface of large assemblies, one shade per chain |
| -membrane | optional | Protein in its lipid bilayer (MD or MemProtMD systems) |
| -electrostatic | optional | Surface colored by Coulomb potential, with color bar |
| -lipophilic | optional | Surface colored by lipophilicity potential, with color bar |
| -property | optional | Cartoon colored by B-factor or user values (e.g. conservation) |
| -density | ligand, peptide | Site with $2F_o - F_c$ electron-density mesh |
| -superpose | optional | Two superposed structures, RMSD reported |

### Targeted protein degradation

Proteolysis-Targeting Chimeras (PROTACs) and molecular glues promote interactions between proteins, enabling targeted degradation through the ubiquitin–proteasome system. In May 2026, vepdegestrant, an estrogen receptor *α* (ER*α*) degrader, became the first FDA-approved PROTAC [15]. Ternary complexes pose a challenge for molecular visualization because the degrader bridges two proteins. Figure 3 illustrates two examples: a 14-3-3-directed molecular glue-based PROTAC recruiting VHL to ER*α* [16], shown as a complete assembly (a) and a close-up of the degrader-mediated interface (b); and the clinical SMARCA2 degrader PRT3789 [17], shown with interface residues colored by protein (c) and as a surface representation of the induced binding pocket (d). Together, these views reveal how degraders engage multiple proteins and organize their interaction interfaces.

**Figure 3:**
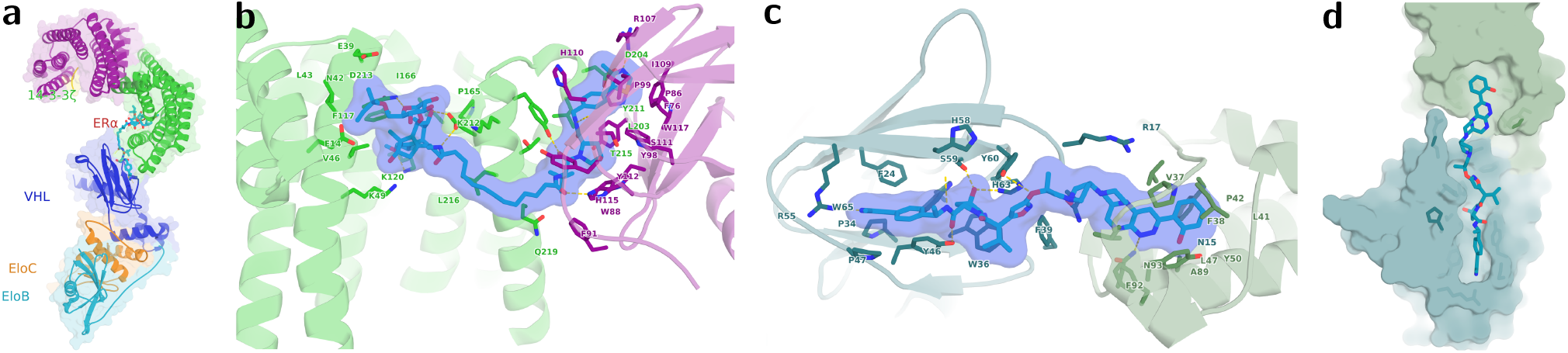
Targeted protein degradation. (**a, b**) A 14-3-3-directed molecular glue-PROTAC recruiting VHL–EloB–EloC to ER*α* (pdb `9SV3` [16]): assembly overview (a) and the degrader bridging 14-3-3*ζ* (green) and VHL (magenta) (b, `–site`). (**c, d**) The SMARCA2 degrader PRT3789 in a ternary complex with the SMARCA2 bromodomain and VHL (pdb 9DTY [17]): binding site with residues colored by protein (c, `–site`) and the degrader lying across the induced interface (d, `–pocket`).

### Emerging viruses

Highly pathogenic avian influenza A(H5N1) viruses of clade 2.3.4.4b have spread to dairy cattle and caused human infections. Figure 4 illustrates two H5N1 hemagglutinin (HA) complexes: HA from A/Texas/37/2024 bound to the human monoclonal antibody H91 [18], shown as a complete assembly (Fig. 4a) and an antibody–antigen interface in which PROTFIG automatically recognizes the Fab as a two-chain protein partner (Fig. 4b); and HA from a cattle-infecting H5N1 virus bound to a human-type receptor analog [19], highlighting binding-site residues and hydrogen bonds (Fig. 4c) and the local electrostatic environment (Fig. 4d).

**Figure 4:**
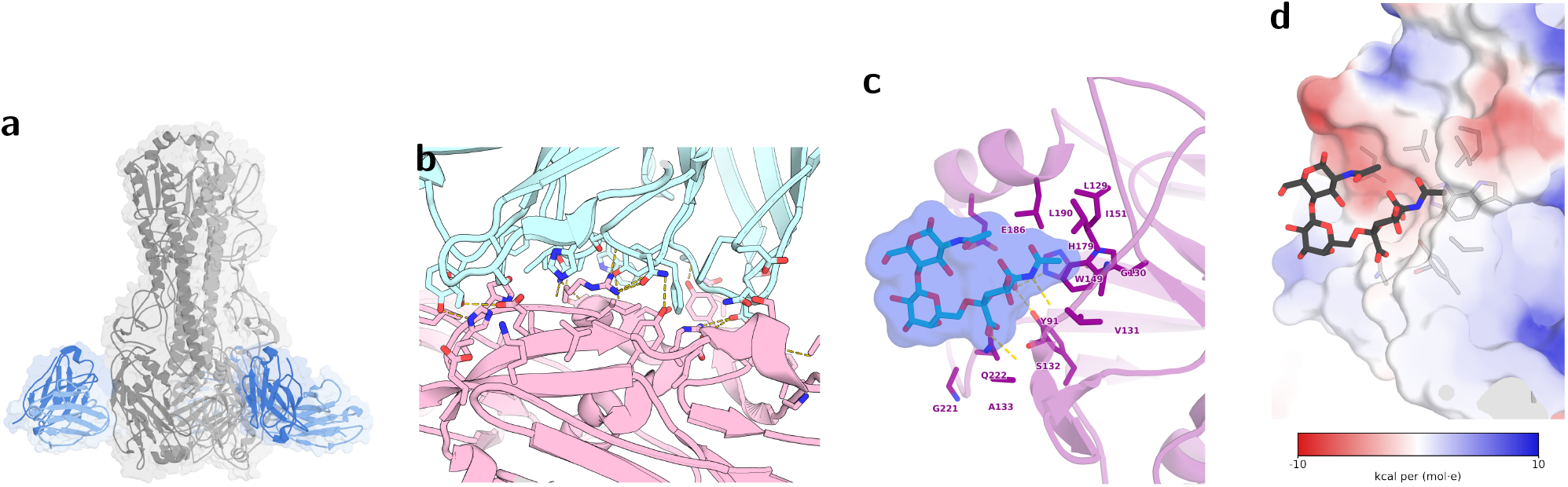
Hemagglutinin of H5N1 clade 2.3.4.4b. (**a**) HA trimer of A/Texas/37/2024 (grey) bound to three copies of Fab H91 (blue; heavy and light chains in dark and light blue) (pdb `9ZTJ` [18]; `–overview` with per-chain colors). (**b**) The H91–HA interface, with the Fab automatically treated as a two-chain protein partner (`–interface_closeup`). (**c, d**) Receptor-binding site of a cattle-infecting H5N1 HA bound to a human-type receptor analog (`pdb` 9JN0 [19]): site residues and hydrogen bonds (c, `–site`) and electrostatic environment of the site (d, `–electrostatic –region site`).

### Other applications

Beyond the examples presented here, PROTFIG has also been used to prepare molecular figures for two recent research articles: one focused on the curation and optimization of protein–peptide structural datasets [20], and another investigating the MxA–Thogotovirus nucleoprotein interface underlying viral restriction and escape [21]. These applications illustrate its utility across different structural biology research workflows.

### Performance

The eighteen panels of this article (Figures 1–4; `–figures.toml` in the repository) were rendered at 300 dpi in 4 min 47 s on a laptop with 11 CPU cores using eight parallel jobs, and as drafts in 27 s.

## 4 Discussion

PROTFIG shifts molecular figure preparation from manual viewer manipulation to style selection. For experimentalists, it enables consistent panels across multiple systems, including variants, ligands, and time points. For structural biologists, saved PyMOL sessions and selections preserve the flexibility to refine each representation. For authors and readers, recorded commands and settings support reproducibility. Because presets are implemented as compact Python functions built on shared components for partner selection, camera orientation, contacts, and labeling, new styles can be introduced with minimal additional code.

Several limitations remain. Hydrogen bonds are identified using geometric criteria rather than energetic calculations and may require manual inspection. Electrostatic and lipophilicity surfaces provide qualitative approximations rather than quantitative predictions. Although automatic camera orientations work well for the examples presented, they may require refinement through the saved PyMOL sessions. Finally, PROTFIG generates individual panels rather than assembled multi-panel figures, leaving their arrangement to the authors; editable label layers and SVG color bars facilitate this process. Future developments will focus on automated multi-panel layouts and AlphaFold-specific presets that visualize model confidence.

## Data and code availability

PROTFIG is open source and available at https://github.com/AmericaChi/PROTFIG under the MIT license. It requires PyMOL (open source or commercial) with Python ≥3.11, NumPy, Pillow and Matplotlib. All structures used are public PDB entries (6XOX, 7KI0, 8YW3, 9SV3, 9DTY, 9ZTJ, 9JN0) and a MemProtMD simulation of 6XOX. The file `–figures.toml` that regenerates every panel of this article, together with the input-download script, is included in the repository.

## Conflict of interest

The authors declare no competing interests.

## Notes

https://github.com/AmericaChi/PROTFIG

### Competing Interest Statement

The authors have declared no competing interest.

